# Neonatal endotoxin stimulation was associated with long- term innate immune markers and an anti-allergic response in bronchiolar epithelium in spite of allergen challenge

**DOI:** 10.1101/854604

**Authors:** Luciana N García, Carolina Leimgruber, Juan P. Nicola, Amado A Quintar, Cristina A Maldonado

**Affiliations:** Centro de Microscopía Electrónica, Facultad de Ciencias Médicas, Universidad Nacional de Córdoba.; Instituto de Investigaciones en Ciencias de la Salud (INICSA), Consejo Nacional de Investigaciones Científicas y Técnicas (CONICET), Córdoba, Argentina; Departamento de Bioquímica Clínica, Facultad de Ciencias Químicas, Universidad Nacional de Córdoba.; Centro de Investigaciones en Bioquímica Clínica e Inmunología (CIBICI), Consejo Nacional de Investigaciones Científicas y Técnicas (CONICET), Córdoba, Argentina

**Keywords:** Club cell, asthma, pulmonary host defense, antiallergic proteins

## Abstract

Asthma is a heterogeneous disease underlying different medical processes, being the allergic asthma, with an early-onset in childhood, the most common type. In this phenotype, the continuous exposure to allergens produces a Th2-driven airway remodeling process that leads to symptoms and pathophysiological changes in asthma. Strategies as the avoidance of aeroallergen exposure in early life have been tested to prevent asthma, without a clear success. Alongside, several mouse models of aeroallergen challenge have dissected potential homeostatic responses by which environmental microbial stimulation reduces the subsequent allergic inflammation in the offspring. This suggests the onset of underlying preventive mechanisms in the beginning of asthma that have not been fully recognized. In this study, we aimed to evaluate if neonatal LPS-induced stimulus in epithelial host defenses could contribute to the prevent asthma in adult Balb/c mice. For this purpose, we studied the response of bronchiolar club cells (CC) that are situated in the crossroads of the host defense and allergic inflammation, and express specific pro and antiallergic proteins. LPS stimulus in the neonatal life intensified the production of TLR-4, TNFα, and natural anti-allergic products (CCSP and SPD), changes that contributed to prevent asthma triggering in adulthood. At epithelial level, CC skipped the mucous metaplasia, declining the overproduction of mucin via the EGFR pathway and the mice expressed normal breathing patterns in front of OVA challenge. Furthermore, the overexpression of TSLP, an epithelial pro-Th2 cytokine was blunted and normal TSLP and IL-4 levels were found in bronchoalveolar lavage (BAL). Complementing this shift, we also detected lower eosinophilia in BAL while an increase in phagocytes as well as in regulatory cells (CD4+CD25+FOXP3+ and CD4+IL-10+) was seen, whit an elevation in IL-12 and TNFα secretion. Summarizing, our study pointed to stable asthma-preventive effects promoted by neonatal LPS-stimulation; the main finding was the increase of several anti-Th2 specific proteins at epithelial level, together with an important diminution of pro-Th2 TSLP, conditions that promoted changes in the local immune response with Treg. We thus evidenced several anti-allergic dynamic mechanisms overlying in the epithelium that could be favored in an adequate epidemiological environment

## INTRODUCTION

Asthma is a heterogeneous disease with diverse underlying processes and many clinical expressions. The most common phenotype is the allergic asthma that has an early-onset in childhood. This phenotype is associated with a family history of allergic diseases, and is characterized by chronic airways inflammation, with activated mast cells, increased numbers of eosinophils, T cells, natural killer T cells, and CD4+ T helper (Th) 2 cells that release IL-4, IL-13, and IL-5. Additionally, IgE-secreting B cells are induced during the asthma process (1, 2). In this phenotype, the continuous exposure to allergens produces several consequences in the structure and function of the airways, with the establishment of a remodeling process that includes mucus hypersecretion, smooth muscle hyperplasia, subepithelial fibrosis, blood vessel proliferation and the infiltration of inflammatory cells (3). All these effects provoke airways narrowing, a common final pathway to symptoms and physiological changes in asthma (1). However, the avoidance of airborne allergen exposure in the early life has been tested in randomized clinical trials and has not been successful in preventing asthma development, suggesting underlying mechanisms in the beginning of asthma that have not been fully recognized so far (4, 5).

The progressive rise in allergic diseases in the last decades denotes the involvement of environmental factors in their pathophysiology (6). Based on epidemiological evidence, the hygiene hypothesis (HH) infers that the reduction of the early life infections due to the modern lifestyle weakens their protective effects against allergic disorders (7). In correlation, recent studies showed low childhood prevalence of allergy/asthma in rural areas as compared with urban areas, related to the perinatal microbial exposure, mainly to the high levels of endotoxins present in dust samples (8–12). In addition, several mouse models have dissected potential immune mechanisms by which environmental microbial stimulation, including the perinatal lipopolysaccharide (LPS) stimulus, of the airways mucosa reduces the allergic inflammation to airborne allergen challenge in the offspring, while favoring homeostatic responses (13–22).

In steady state, the homeostasis of the airways relays on the bronchioalveolar cells (23, 24). For this purpose, airways epithelial cells (AECs) express inflammatory, anti-inflammatory, chemoattractant, antimicrobial mediators, as well as pattern recognition receptors (PRRs) to detect environmental molecules as endotoxin, and initiate an innate immune response by activating dendritic cells (DC) (25). This link between innate and adaptive immunity has evidenced a significant role of AECs in lung immunity and highlighted that an abnormal epithelial response may lead to a chronic inflammatory response (26).

When AECs take contact with inhaled stimuli, which contain multiple proteolytic allergens as well as microbial contaminant, they are induced to produce ROS and pro-Th2 cytokines like TSLP, IL-25 and IL-33. These cytokines interact by cell–cell communications with subepithelial DC, mast cells as well as innate lymphoid cells, which in turn trigger the recruitment of Th 2 cells, leading to an amplified Th-2 cytokines production in the airways (27, 28) (29) (24, 30).

Additionally, there is accumulative evidence about AECs intrinsic alterations in childhood asthma that render airways more vulnerable to airborne allergens and predispose them to Th2- responses (31–34). These data indicate that AECs are essential controller of the immune response to allergens and may be an early player in order to bias a Th2 response in the immature immunity system. Therefore, AECs play a particular role since they are situated at the crossroad of the innate host defense and allergic inflammation.

Such contrasting activities are clearly exemplified by bronchiolar club cells (CC). They perform a myriad of homeostatic mechanisms including detoxification of xenobiotics and being a stem/progenitor cell to the airways AECs (35, 36). Additionally, CC directly contribute to host defenses by secreting monocyte and neutrophil chemoattractants, the antibacterial collectin surfactant protein (SP) D and the anti-inflammatory club cell secretory protein (CCSP) (37–43). However, under allergic genetic predisposition, CC can also activate elicited a Th2-inflammation via IL-4 receptor, driving eosinophil accumulation by producing eotaxin. Furthermore, they are the principal cells to undergo epidermal growth factor receptor (EGFR)-mediated mucous metaplasia as demonstrated in experimental models of asthma (29, 44–46). Interestingly, both SP-D and CCSP, play a direct role suppressing allergic inflammation in vivo and in vitro, inciting Th1 cytokines increment under LPS-stimulus. There is quite evidence of the reduction of these mediators in allergic/asthmatic patients as well as in mouse models of asthma (37, 40, 41, 47–54).

The potential of CC to respond to Th1 inflammatory stimulus, activating protective mechanisms, has often been applied in studies to evidence if this protective role of epithelium prevents the development of Th2 inflammation. In a previous study we reported that LPS pre-exposition to the allergen sensitization partially avoids mucous metaplasia of CC. In consequence, the loss of anti-allergic products in CC and alveolar adult mice macrophages were prevented. We observed a reduction of eosinophil influx, Interleukin-4 levels and airway hyperreactivity, while the T-helper type 1 related cytokines IL-12 and Interferon-g were enhanced (55). Considering early life as a better window of opportunity for triggering an appropriate maturation of innate immunity. In this study, we aimed to evaluate if LPS-stimulation during the neonatal lapse provides better asthma-preventive effects to preserve adult AECs from the Th2-driven inflammation. Mainly evaluating the role of bronchiolar CC and the preservation of their pro and antiallergic proteins.

## MATERIALS AND METHODS

### Animals

Balb/c mice were provided by Fun Vet (Universidad Nacional de La Plata, Argentina) and housed under controlled temperature and lighting conditions, with free access to tap water and commercial lab chow (GEPSA FEEDS, Buenos Aires, Argentina). Animals were randomly assigned to four groups (n= 6 each) and experiments were repeated at least three times.

The animal care and experiments were conducted following the recommendations of Helsinki convention, and in compliance with local laws on the ethical use of experimental animals.

### Experimental design

#### Neonatal treatment

Offspring Balb/c mice were exposed to intranasal applications on every second day from days 3 to 13 of life. While one group of animals were sham treated with PBS, the other received LPS (1μg/5μl; *Escherichia coli* O55:B5 Sigma-Aldrich; St. Louis, MO, USA) according to protocols previous optimized for volume (16) and treatment timing (19).

#### Allergen sensitization

At the age of 4 weeks, all female animals were selected and sensitized by subsequent i.p injections of 0.1 ml of OVA grade VI (1000 μg, Sigma-Aldrich) absorbed to 1 mg of imject Alum (Pierce Rockford, USA) on the 4 and 6 weeks of life.

#### Airway challenge

Ten days later, neonatally (n) LPS-treated mice as well as PBS-exposed were divided into 2 groups. Whereas LPSn/OVA and PBSn/OVA mice were challenged daily (on 10 consecutive days) by an intranasal application of 50μl of 1% OVA, LPSn and PBSn mice were submitted to intranasal application of saline (Fig 1a). Then, after 24h, mice were sacrificed and processed according to the specific methods outlined below.

**Fig 1.**
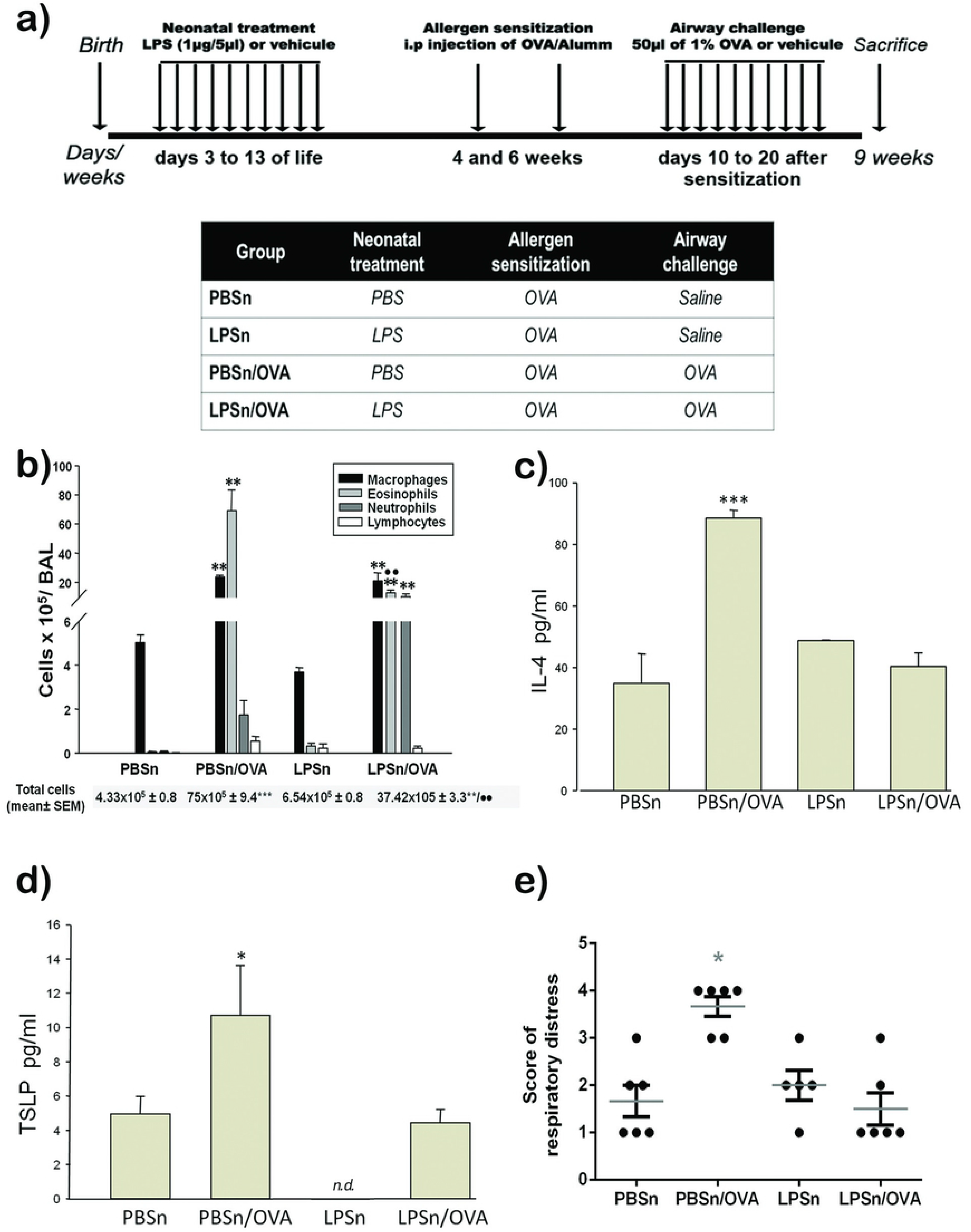
Experimental design and allergic inflammatory state. (a) Timeline diagram and protocols employed in this study. (b) Differential quantification of cell populations in bronchioalveolar lavage (BAL). Bar graph represent total number of macrophages, esosinophils, neutrophils and lymphocytes in BAL. (c) and (d): Levels of IL-4 and TSLP by ELISA. (e) Score of respiratory distress. The score represents increasing signs of respiratory distress obtained in the first minute after intranasal challenge in all groups. Data represent mean ± SD ***p<0.001 vs PBSn,** p<0.01 vs PBSn, *p<0.05 vs PBSn, ●● p <0.01 vs PBSn/OVA

### Lung histopathology

Morphological analysis was conducted in the right lungs of 3 mice per group as previously described. Briefly, at least in 3 experiments, lungs were differentially fixed for either ultrastructural or histopathology analysis by intratracheal perfusion and processed in order to being examined under electron microscope (Zeiss LEO 906E) or light microscope (Axiostar Plus, Zeiss, Germany).

### Mucous cell staining

The Alcian blue-periodic acid Schiff (AB-PAS) staining technique as previously described identified Mucous-secreting cells in the bronchiolar epithelium. Photomicrographs at ×400 were taken using a light microscope equipped with a digital camera (Axiocam ERc5s). A total of 20-30 bronchioles (900-1700 μm diameter) per mouse were analyzed, and the numbers of AB-PAS positive cells present in epithelia lining per 100μm of basement membrane were quantified using Image J Software (NIH version 1.43).

### Immunohistochemical analysis of lung tissue

Immunohistochemical staining was performed as described elsewhere. Briefly, after being blocked, the sections were incubated overnight at 4°C with antibodies recognizing SP-D (1:1000-Chemicon, Temecula, CA, USA), TNFα (1:50 - Hycult, Plymouth Meeting, USA), CCSP (CC10 antibody 1:1000 - Santa Cruz Biotechnology, Santa Cruz, CA, USA), TLR4 (1:100- Santa Cruz Biotechnology), TSLP (1:200- Gene Tex, USA) or pEGFR (1:50 - Santa Cruz Biotechnology), with bound antibodies being detected using anti-rabitt (for SP-D, TNFα, TSLP and CCSP) or anti-goat (for TLR4 and pEGFR) biotin-labeled antibodies (Vector Laboratories, Burlingame, CA, USA) in 1% PBS-BSA. The sections were then incubated with ABC complex (VECTASTAIN Vector Labs, Southfield, MI, USA). Diaminobenzidinde (DAB, Sigma-Aldrich), which was used as a chromogen substrate, and the bronchioles (700-1400 μm diameter) were analyzed and photomicrographs × 400 were taken.

### Bronchoalveolar lavage collection and cell counting

Bronchoalveolar lavage (BAL) were obtained (n= 9 mice/group in three different experiment) as described elsewhere (52). Briefly, after three serial intra-tracheal instillations of 1 ml PBS, the cells obtained were centrifuged at 200g, resuspended and counted meanwhile the supernatant was stored at −70°C for ELISA.

For cytospin preparations, about 12.5×10^4^ cells from the pellets were cytocentrifuged onto slide, whereas some slide were preserved at −70°C for immunofluorescence, others were stained with May Grünwald-Giemsa (Biopur Diagnostic, Rosario, Argentina) and counted. The cell populations were evaluated for two samples per mouse, and a total of 2400 cells per group were counted.

### Immunofluorescence

Cytospin preparations (3 per mice) obtained from the BAL (3 mice per group) were withdraw at room temperature at immediately fixed with 4% formaldehyde, permeabilized with 0.25% Triton X-100 in PBS and incubated for 1 h in 5% PBS-BSA to block non-specific binding. Slides were double immunostained by incubating overnight at 4°C with a mix of anti-CD4 conjugated with PERCP (BioLegend, San Diego, CA, USA) and anti IL-10 conjugated with PE (BD Biosciences Pharmingen, San Diego, CA) and mounted using fluoromount containing DAPI. Afterwards, the cells were viewed with Fluoview 1000 Confocal and laser scanning microscope, (Olympus, Tokyo, Japan) and serial × 60 microphotographs (10 per coverslide) were collected, with all double immunostained cells being evaluated in three different experiments and the relative percentages were calculated.

### Flow cytometry

Pellet cells obtained from BAL (n= 5 mice/group in three different experiment) were incubated with a mix of conjugated antibodies (Biolegend) for the following T-cell subset superficial markers: APC-Cy7 anti-mouse CD45 (1:600); FITC anti-mouse CD4 (1:200); PerCP anti- mouse CD25 (1:200) for 30 min at 4°C. Next, the cells were fixed (CITOFIX; BD Biosciences Pharmingen, San Diego, CA) for 20 min at 4°C and permeabilized with Perm/Wash (BD Biosciences Pharmigen), before being incubated with a dilution 1:30 of the intracytoplasmic antibody: APC anti mouse FOXP3 (eBIoscience) for 30 min at 4°C. Finally, the cells were washed, suspended in filtered PBS (1×10^5^ events/experimental treatment), and analyzed by flow cytometry (FACSCanto II Flow Cytometer, BD Biosciences, San Diego, CA, USA). Data analysis was carried out using the FlowJo software (Tree Star, Ashland, OR).

### Immunobloting

By Western Blot SP-D, TLR4 and TNFα levels were determined in total lung homogenates from 3 animals per group in three different experiments as was described (52). Briefly, after proteins were measured with a Bio-Rad kit (Bio-Rad Laboratories, Hercules, CA, USA), the denatured protein samples were separated on 12% SDS-PAGE and blotted onto a Hybond-C membrane (Amersham Pharmacia-GE, Piscataway, NJ, USA). Membranes were then blocked with 5% defatted dry milk in TBS/0.1% Tween 20, and incubated for 3h with one of the following antibodies: rabbit anti-SP-D (1:1000 - Chemicon, rabitt anti TNFα (1:50 – Hycult) or mouse anti-TLR4 (1:300 Abcam, Maryland, USA). Blots were incubated with a peroxidase-conjugated (HRP) anti-rabbit (Jackson Immunoresearch Labs Inc, West Grove, PA, USA), or anti-mouse (Jackson Immunoresearch) secondary antibodies at a 1:2000 dilution. Finally, the membranes were rinsed in TBS/0.1% Tween-20 and exposed to Pierce™ ECL Western Blotting Substrate (Thermo Fischer Scientific) following the manufacturer’s instructions. Emitted light was captured on Hyperfilm (Amersham-Pharmacia) and a densitometry analysis was performed by applying the Scion Image software (V. beta 4.0.2, Scion Image Corp., Frederick, MD, USA). Additionally, the expression of ACTB (1: 4:000; mouse anti-βactin; Sigma-Aldrich) was used as an internal control to confirm equivalent total protein loading.

### Dot Blot Analysis

The CCSP protein expression was evaluated in lung homogenates after protein measurement was performed using a Bio-Rad kit. Samples were then adjusted to 5μg/μl in PBS, pH 7.4, and 10μl of each sample were spotted onto a Hybond C membrane (Amersham Pharmacia). Then, the membrane was blocked with 5% fat-free milk in PBS buffer for 1h and then incubated for 3h with a rabbit primary antibody anti-CC10 1:500 (Santa Cruz Biotechnology) in blocking buffer at room temperature. After washing with TBS–Tween-20 buffer, the membrane was treated with a HRP-conjugated anti-rabbit antibody (Jackson Immunoresearch) and the next handle was as described above for Western blot.

### Cytokine detection by ELISA

Cytokines production was measured, in BAL supernatant, following the manufacturer’s instructions. It was applying commercially available sandwich ELISA kits for IL-4 (BD Biosciences), IL-12 and TSLP (Biolegend, San Diego, CA, USA), as well as TNFα and IFNγ (eBioscience, San Diego, CA, USA).

### RNA isolation and gene expression analysis

Total RNA was extracted from right lung tissue samples (~0,01mg) with Trizol reagent. RNA was subsequently purified using Direct-zol RNA min prep kit (Zymo Research) following the manufacturer’s instructions and quantified with a ND-1,000, NanoDrop spectrophotometer (Thermo Scientific) at 260 nm. Measurements of A260/280 were used to determine the purity of the RNA. After that, 1μg of RNA was used as template for reverse transcription following the manufacturer’s instructions (EpiScript™ Reverse Transcriptase System kit, Epicentre, USA), using random hexamer primers (Fermentas, Thermo Fisher Scientific, MA, USA) and utilizing a My Cicle rTM BIO-RAD (Thermal Cycler System, CA, USA).

Real-Time PCR analysis was performed on an ABI Prism 7500 detection system (Applied Biosystem, CA, USA) using Power SYBR Green PCR Master Mix (Applied Biosystems, Thermo Fisher Scientific). Relative changes in gene expression were calculated using the 2-ΔΔCt method normalized against the housekeeping gene 18s. For each pair of primers, a dissociation plot resulted in a single peak, indicating that only one cDNA species was amplified. Amplification efficiency for each pair of primers was calculated using standard curves generated by serial dilutions of cDNA. All primers were from Invitrogen (Buenos Aires, Argentina). The specific primers pairs used were: TSLPfp: 5’-AGAGAAATGACGGTACTCAGG-3’, TSLPrp: 5’-TTCTGGAGATTGCATGAAGGA-3’; 18sfp 5’-ATGCGGCGGCGTTATTCC-3’, 18srp: 5’-GCTATCAATCTGTCAATCCTGTCC-3’; CCSPfp: 5’-GATCGCCATCACAATCACTG-3’, CCSPrp: 5’-CTCTTGTGGGAGGGTATCCA-3’; SP-Dfp: 5’-TGGACCCAAAGGAGAGAATG-3’, SP-Drp: 5’-CATGCCAGGAGCACCTACTT-3’.

### Clinical score assessment of the degree of respiratory distress

In three different protocols, at days 7-10 of the allergen challenge, the breathing patterns of mice (n=6/group) were video recording during the first minute after OVA instillation. The values assigned to increasing signs of respiratory distress resulted from the adaptation of the respiratory failure clinical score system developed by Wood (56). The scoring was performed, via a double-blind procedure, by three different physician operators and analyzed by a nonparametric statistical test (see below).

### Statistical analysis

In general, data obtained were analyzed by one-way ANOVA, followed by post-hoc comparison with the Tukey-Kramer test. In particular, for the analysis of clinical score data, we applied the Kruskal Wallis test. For all test a p<0.05 significance level was used.

## RESULTS

### Neonatal LPS stimulation inhibited OVA-induced allergic airways inflammation triggered in the adulthood

Firs, we evaluated the effect of an early exposition with LPS (represented in the LPSn/OVA group) over an ulterior OVA allergic response in the airway’s analyzing BAL cytokines and cellular inflammatory content as well as the breathing pattern recorded in the different mice groups.

We detected that neonatal LPS treatment affected the development of experimental asthma triggered in adulthood. As shown in Fig.1, the establishment of experimental asthma in adult mice was largely prevented as indicated by a significantly lower influx of both total inflammatory cells and eosinophils into the airways lumen of LPSn/OVA mice compared to the PBSn/OVA group (Fig.1b). In addition, the number of macrophages remained unchanged while neutrophils increased significantly (Fig 1a). Surprisingly, IL-4 and TSLP, both associated to Th2 inflammation, exhibited normal levels in BAL of LPS pre-treated mice in spite of the allergen-challenge, while they were significant higher in PBSn/OVA group (Fig. 1c and 1d respectively). As expected, in the LPSn group neither BAL cell count nor IL-4 content were different from controls (Fig 1b and c); TSLP was remarkable reduced to non-detectable levels (Fig. 1d).

To test whether the inflammatory parameters were accompanied by changes in the degree of respiratory distress, a clinical scoring system was carried out (See Supplementary material). While most of the neonatal PBS-exposed mice displayed higher signs of respiratory distress after OVA challenge, the breathing pattern of LPSn/OVA mice was not different from control mice (Fig 1e).

### Neonatal LPS application promoted innate immunity mediators, antimicrobial cytokines and Treg cells in the airway’s microenvironment

After demonstrating the abrogation of a Th2 inflammatory response, we investigated whether neonatal LPS exposure influences other components of the immune response. In the airway’s milieu LPS stimulus increased inflammatory cytokines as TNFα and IL-12 in both LPSn and LPSn/OVA mice groups (Fig. 2c and 2b, respectively). It was noteworthy that LPS induced high levels of IFNγ, a prototypical Th1 cytokine, in the LPSn group but not in LPSn/OVA animals (Fig. 2a). By contrast, the latter exhibited the higher influx of CD25+/FOXP3+ Treg cells (48,16 % ± 10,2 LPSn/OVA vs 19.65 % ± 4,22 PBSn/OVA) (Fig.2d). Furthermore, immunofluorescence performed in cytospins displayed an increased level of IL-10 positive cells in LPSn/OVA group compared with PBSn animals, although the analysis of the IL-10+/CD4+ cells ratio revealed a significant change in both group (LPSn and LPSn/OVA) compared to control. (Fig. 2e).

**Fig 2.**
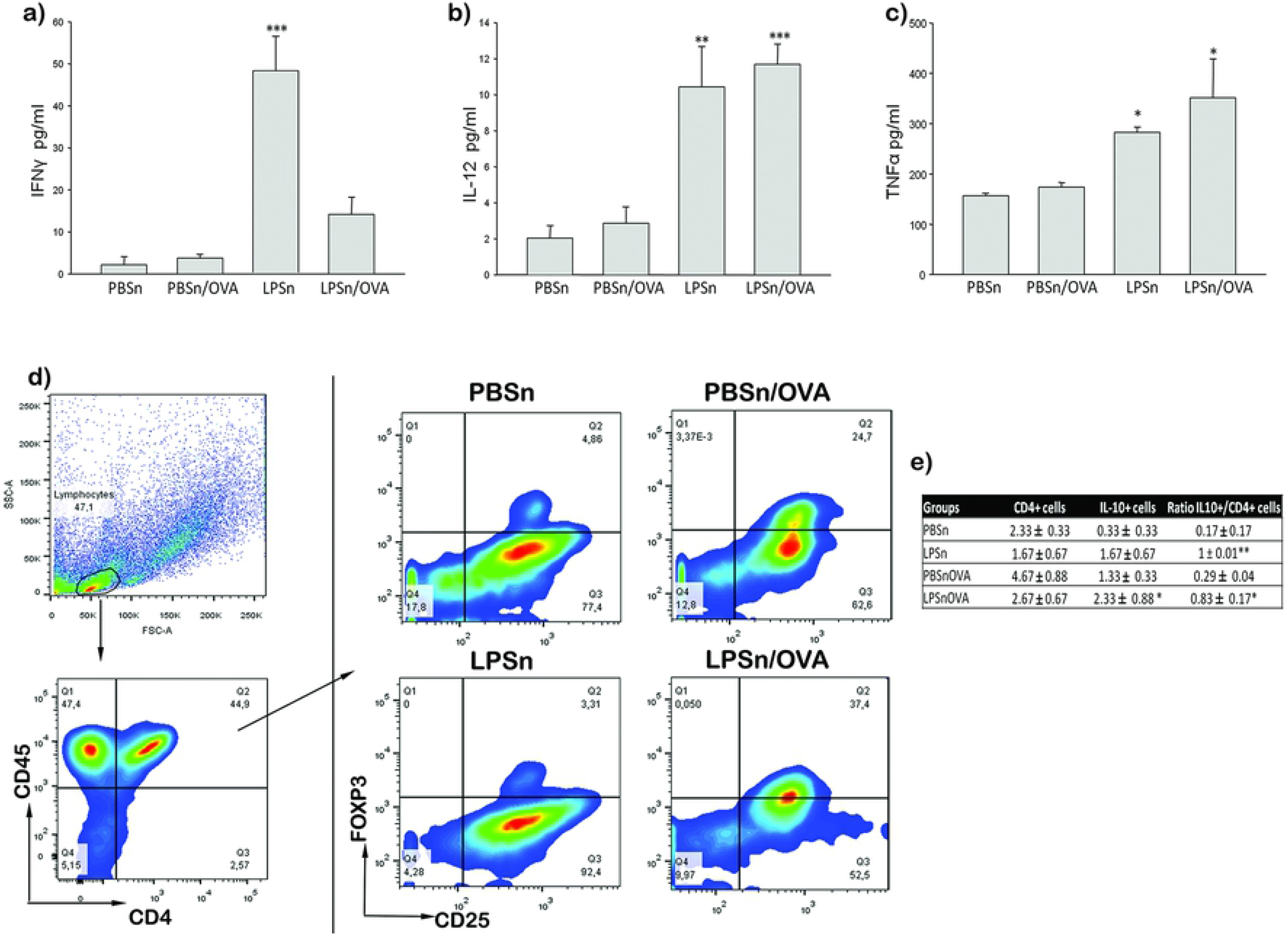
Modulatory response of the airway’s environment in BAL. a), b) and c) IFNγ, IL-12 and TNFα levels by ELISA, respectively. d) Percentage of CD4+CD25+FOXP3+ cells obtained in BAL by flow cytometry. The plots correspond to a representative experiment analysis in all groups, the % of CD25+FOXP3+ cells are shown in the Q2 quadrant. e) Immunofluorescence count of CD4+, IL 10+ and ratio of IL10+/CD4+cells performed in cytospin. Data are represented as mean ±SEM, *p< 0.05 vs PBSn, **p<0.01 vs PBSn, ***p<0.001vs PBSn.

We next characterized the epithelial response in this long-lasting anti-allergic modulation in order to evidence if the early LPS stimulus can trigger local epithelial mechanisms involved in prevention of asthma development in adulthood.

### Neonatal LPS exposure abrogated the development of mucous metaplasia and pro-allergic mediators in the bronchiolar epithelium

We first analyzed changes in the expression of specific pro-allergic mediators that are known to increase in bronchiolar epithelium during asthma,

As was shown before (55), the OVA-allergic inflammation incited mucous cell metaplasia in the bronchiolar Club cell via EGFR signaling. In this way, Figure 3 shows an increased number of mucous secreting cells (AB-PAS panel in Fig 3a and Fig. 3b) as well as the overexpression of phosphorylated-EGFR in the apical cytoplasm of CC in PBSn/OVA mice (pEGFR panel in Fig. 3a). Whereas in LPSn/OVA mice, both, pEGFR overexpression (pEGFR panel in Fig. 3a) and mucous metaplasia (Fig. 3b), were largely reduced by the neonatal endotoxin-treatment.

**Fig 3.**
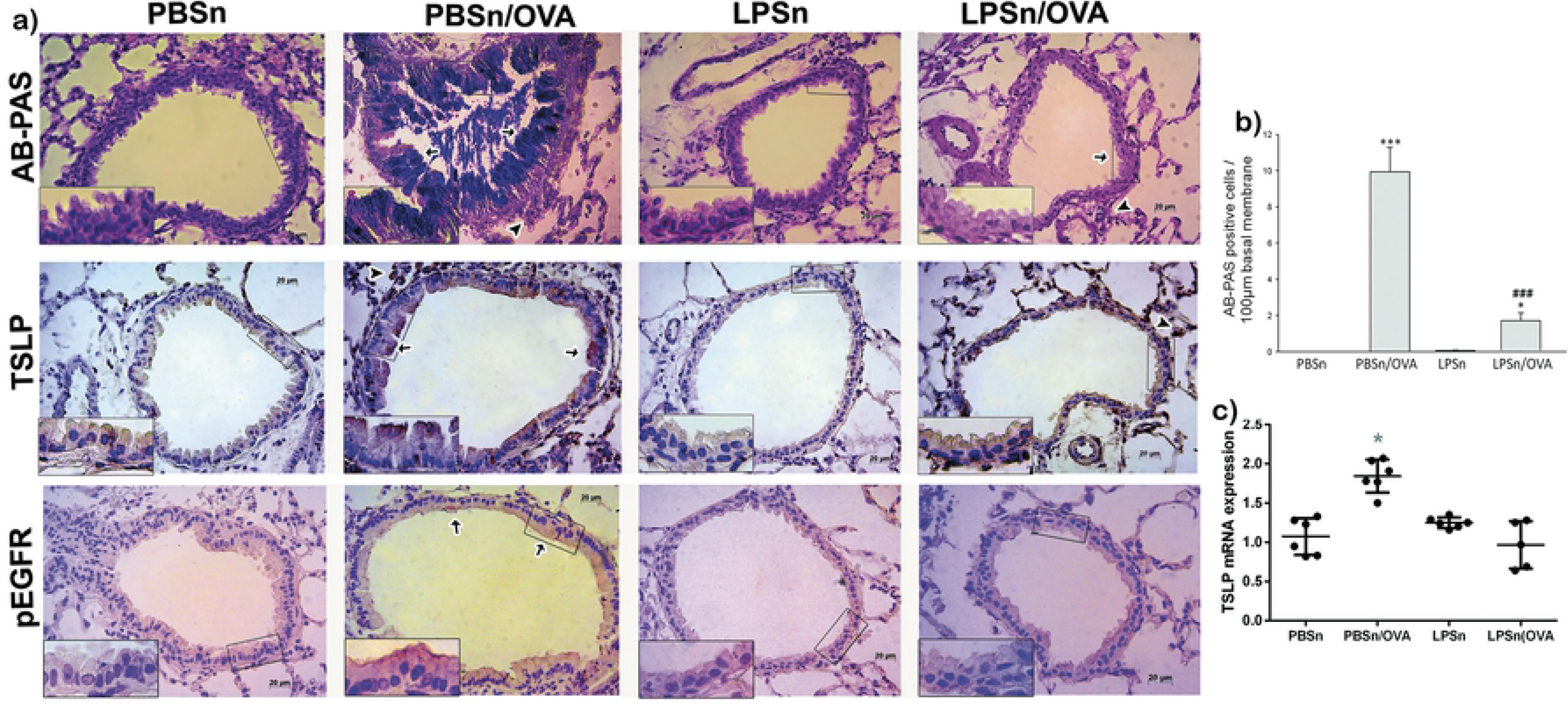
Mucous metaplasia analysis in Club cells and epithelial TSLP expression. a) Representative photomicrographs of Alcian blue-periodic acid Schiff (AB-PAS), TSLP and pEGFR staining of bronchiolar sections. Scale bars: 20μm. In AB-PAS panel arrows indicate AB-PAS positive cells in PBSn/OVA and LPSn/OVA groups, while arrowheads indicate infiltrating inflammatory cells. In TSLP panel, arrows indicate positive cells in PBSn/OVA although some cells (arrowhead) from te inflammatory response also expressed TSLP, inset selection demonstrated the lack of staining in club cells LPSn/OVA groups.. In pEGFR panel arrows indicate positive cells in PBSn/OVA, while the inset demonstrated the apical expression of the activated receptor in club cells. b) Graph represents AB-PAS cells count per 100μm. c) TSLP mRNA expression by Real-Time PCR analysis. Graph represents fold increase expression in lung tissue homogenate. Data are represented as mean ±SEM, *p<0.05 vs PBSn, ***p<0.001 vs PBSn, ### p<0.001 vs PBSn/OVA.

We also studied the effect of neonatal-LPS on the expression of TSLP, an epithelial cell cytokine that promotes Th2 differentiation after allergen contact. In accordance with the pEGFR and mucous metaplasia induction, bronchiolar epithelial cells of PBSn/OVA group, showed strong TSLP immunoreactivity in the apical cytoplasm; meanwhile CC of LPSn/OVA animals skipped of the overexpression of TSLP (TSLP panel in Fig 3a). These results were corroborated by the quantitative PCR analysis (Fig. 3c), showing that lung TSLP mRNA almost duplicated its expression in PBSn/OVA animals (1.85 ± 0.09 PBSn/OVA vs 1 ± 0.1 PBSn) while remained unchanged in LPSn/OVA mice (1.12 ± 0.19).

In previous studies, we have demonstrated that the ultrastructure of CC is a sensitive parameter of the airways allergic inflammatory affectation (Roth 2007, 2013, Garcia 2014). For this reason, we studied CC morphological profile in all groups by electron microscopy (Fig. 4). At this level, we corroborated the preservation of the typical cellular profile in PBSn mice, characterized by the presence of a dome-shape cupola, numerous polymorphic mitochondria in the cytoplasm, along with scarce spherical electron-dense secretory granules under the plasma membrane (Fig. 4a). These parameters could also be seen in LPSn/OVA mice, which differed only by an increase on the number of normal electron-dense granules as well as a mayor development of RER (Fig. 4d). Meanwhile, PBSn/OVA animals displayed characteristic mucous cell metaplasia featured as a hypertrophied cytoplasm filled up with numerous large electron-lucent secretory granules, slim mitochondria and abundant RER (Fig. 4b). In control mice, only exposed to LPS, CC also developed an increased number of electro-dense granules, as was shown by LPSn/OVA animals (Fig. 4c). In this group, the evident diminution of their CC cupola is probably due to the repeated LPS instillation they received in neonatal life.

**Fig 4.**
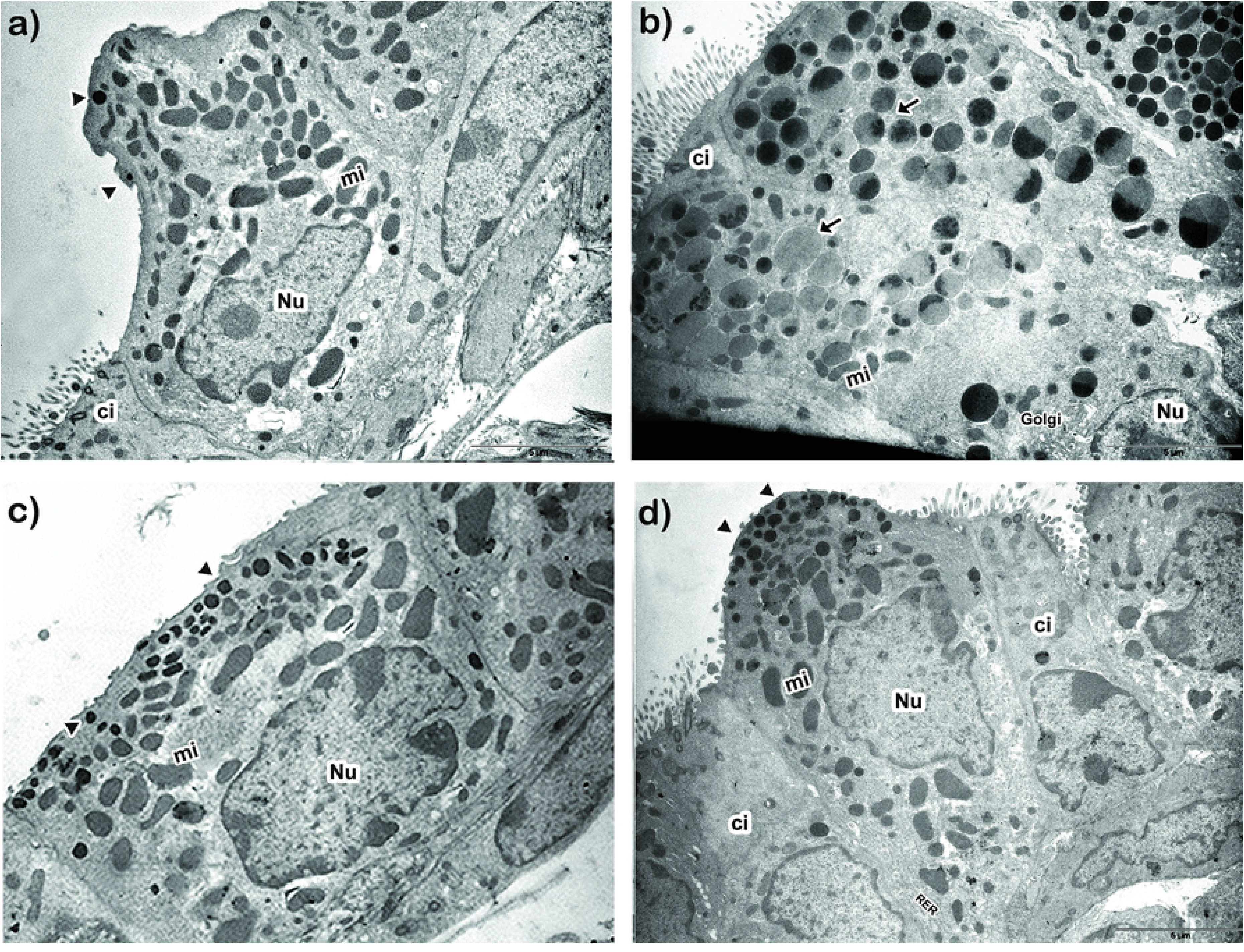
Club cell ultrastructural features. Representative electron micrograph images of club Cell morphology of PBSn (a), PBSn/OVA (b), LPSn/OVA(c) and LPSn/OVA (d) groups are shown. Scale bar represents 5μm. Nu: nucleus, Mi: mitochondria, Ci: ciliated cells, Golgi: Golgi apparatus, RER: rough endoplasmic reticulum. Arrowheads: normal electron dense granules, arrows: electron lucid granules.

### Neonatal LPS stimulus promoted a long-lasting increase of mediators of innate response and Th2-immunomodulatory proteins on bronchiolar epithelium

Next, we analyzed whether the mucous metaplasia prevention by neonatal LPS treatment correlated with changes in the expression of epithelial host defense mediators, mainly CCSP and SP-D. As it was described (55), OVA-allergic inflammation induced a diminution in the imunoreactivity of CCSP and SP-D in CC of PBSn/OVA group when compared to its control group (CCSP and SP-D panels in Fig 5a). Meanwhile for both LPSn and LPSn/OVA groups, a strong CCSP and SP-D immunolabelling was observed (Fig 5a). These changes in protein expression of CCSP and SP-D were also verified by immunoblottting (Fig. 5b and 5c). However, the neonatal LPS-instillation did not increase mRNA expression of CCSP or SP-D in LPSn and LPSn/OVA (Fig. 5f and 5g, respectively). This may be due to the stimulus for protein secretion provided by the allergen challenge in LPS/OVA group, and to the contribution of SP-D of the Type II alveolar cells; which could explain the highest SP-D content by western blot analysis in these group.

**Fig 5.**
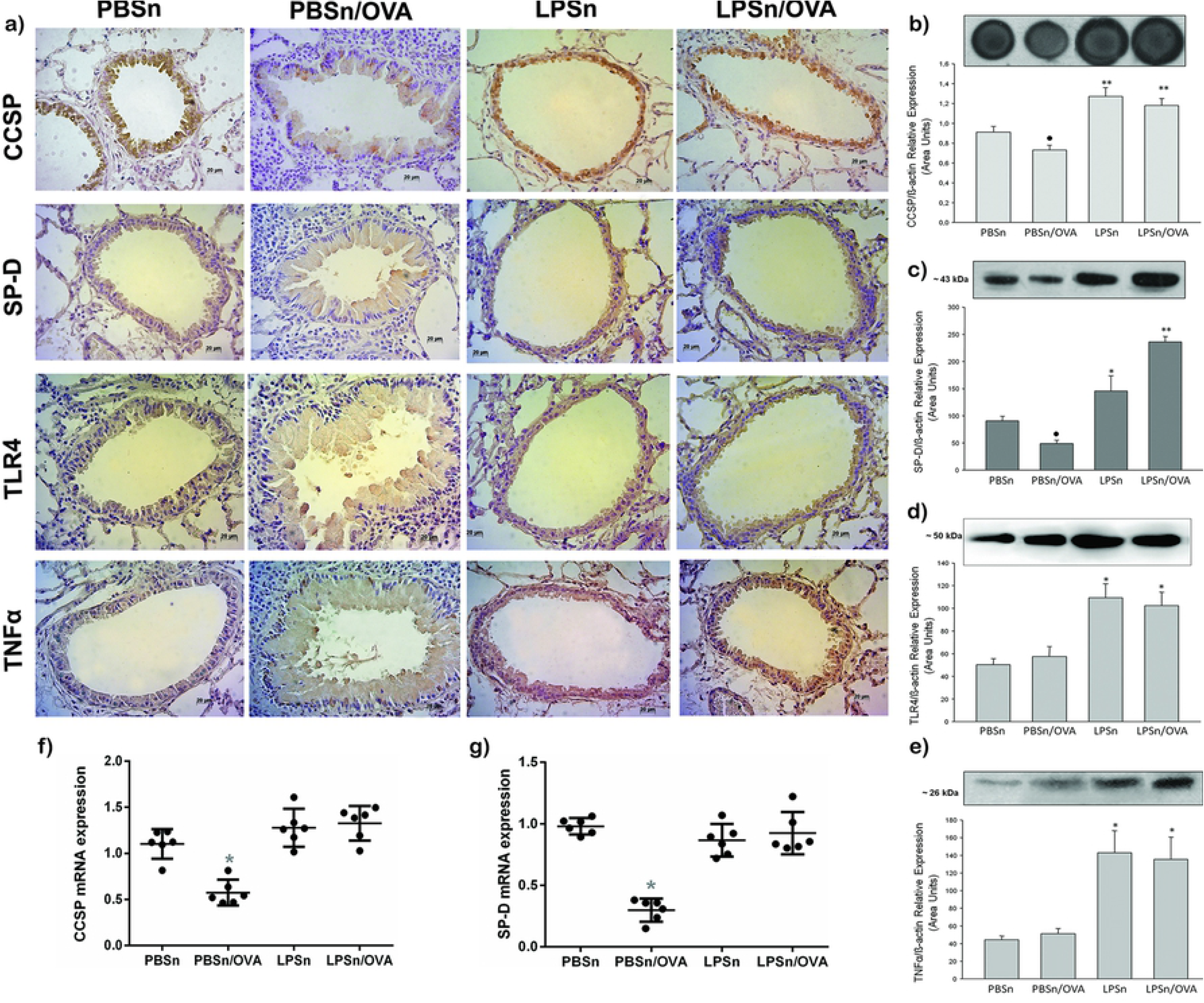
Club cell’s expression of host defense proteins and antimicrobial cytokines. a) Immunostaining of CCSP, SP-D, TLR4 and TNFα performed on lung section of all groups. Positive cells appear in brown against the blue counter stain of haematoxylin. Scale bars: 20μm. b) Dot blot of CCSP in lung homogenates. Graph represents fold increase of the relative CCSP/β-actin expression in lung homogenate by densitometric analysis. c), d) and e) Western Blot of SP-D, TLR4 and TNFα lung content, respectively. Graph represents fold increase of the relative expression in lung homogenate by densitometric analysis.f) and g) CCSP and SP-D mRNA respectively expression by Real-Time PCR analysis. Graph represents fold increase expression in lung tissue homogenate. Data are represented as mean ±SEM, *p< 0.05 vs PBSn, **p<0.01 vs PBSn, ● p<0.05 vs PBSn.

Regarding the microbial recognition and cytokine response, both the toll like receptor 4 (TLR4) and TNFα, increased their expression in CC as well as in lung tissue of both LPS neonatal stimulated groups (Fig 5d and 5e, respectively). These suggest a specific LPS-response in bronchiolar epithelium that induced a persistent elevation of these defense molecules and seemed to be preserved in spite of allergen stimulus.

## DISCUSSION

In the present work, we reported that neonatal LPS-treatment triggers anti-allergic secretory products of the local airway epithelium that persist in adulthood. Among these products, we demonstrated, the increase of CCSP and SP-D content, together with the upregulation of TLR-4 and TNFα, both related to innate host defenses, in epithelial CC and lung tissue. In correlation, CC skipped the mucous metaplasia pathway in front of airborne allergen challenge, preserving their typical phenotype and declining EGFR, mucins, and TSLP, a pro-Th2 cytokine, overexpression. Furthermore, under allergic stimulus, animals with neonatal LPS treatment exhibited normal breathing patterns, normal IL-4 and TSLP levels and a lower eosinophilia. By contrast, in those mice, an increase in phagocytes and in regulatory cells (CD4+CD25+FOXP3+ and CD4+IL-10+), as well as in IL-12 and TNFα levels was observed. These evidences are remarkable considering that they reveal a possible new preventative and therapeutic approach to asthma focused on increasing the airways resistance to environmental insults rather than suppressing Th2 downstream inflammation once it is established. The finding of anti-allergic effects associated to CCSP and SP-D is consistent with previous results (19) (41, 49–51) (40, 48). For instance, there is evidence that these proteins down regulate the type 2 differentiation of Th cells, inhibits the allergen-activation of innate immune cells (eosinophils, basophils, and mast cells) and are reduced in both BAL and serum of asthmatic individuals (40, 41, 50, 51, 57) (47, 48, 58). Moreover, previous reports from our laboratory showed the interplay between the establishment of an experimental asthma model in mice and the diminution of CCSP and SP-D levels, which were restored by Budesonide or Montelukast treatment (52).

Recently, we determined that the pre-treatment of adult mice with LPS before an allergic inflammation partially prevented CCSP and SP-D reduction in CC, and the increase of IL-4 levels and airways hyperresponsiveness (55). However, in the present work we demonstrate that when endotoxin treatment is performed in neonatal life it achieved a more extensive asthma prevention in adulthood. The neonatal treatment not only avoided the metaplastic changes in these cells, but also preserved the mRNA levels of CCSP and SP-D; moreover, the characteristic increment of IL-4 and respiratory distress in front of OVA challenge were damped. These results are similar to the blunting of a Th-2 allergic response and airways hiperresponsiveness (AHR) reported by other authors using either, infant or pregnant mice and microbial stimulus (16, 17, 19) (13)

As expected, an increase in TNFα and IL-12 was observed in both groups exposed to LPS; nevertheless, a robust Th1 response was only seen in LPS exposure animals as indicated by the high IFNγ levels meanwhile in LPSn/OVA the most important immunological change was the increased number of Treg found. In addition, whereas LPSn/OVA group was the only one that reached a significant number of CD4+IL10+ cells, both groups (LPSn and LPSn/OVA) demonstrated a significant ratio of IL10+/CD4+ cells ex vivo versus PBSn animals. Although the experimental design of our study cannot explain the IL-12 elevation coexisting with a Treg response in LPSn/OVA group, other authors have related the persistent increment of IL-12 cytokine as a stimulus of phagocyte activity (59, 60).

Meanwhile studies conducted by Gerhold in adults Balb/c with systemic administration of an anti- IL-12 before LPS stimulus, demonstrated that the reduction of an ulterior allergic inflammation occurs in an IL-12 dependent way (17). Regarding the Treg response, Nguyen et al previously described that TSLP directly impairs the function of pulmonary Treg cells obtained of allergic asthma patients (61). This was indicated by a significant decrease in suppressive activity and IL-10 production compared to healthy control and non-allergic asthmatic counterparts, which were associated with the TSLP expressions levels in BAL. Therefore, it is probable that the diminution of TSLP induced by LPS-pretreatment in this study, had the additional effect of restoring Treg.

In accordance with our results, experimental studies conducted by other authors in neonatal Balb/c mice exposed to LPS and different models of sensitization and exposition to OVA,also evidenced the occurrence of a response involving the expression of IL-10 and IFN-γ in the re-exposition to allergen (16) (19). Furthermore, Gerhold demonstrated that LPS, either in prenatal or postnatal stimulus, induces a persistent elevation in soluble factors such as CD14 and Lipopolysaccharide binding protein-LBP, as well as TLR4 mRNA expression in young mice (16). More recently, the gene expression levels of innate and adaptive immunity essential markers in white blood cells in farmers’ children were assessed in the multinational and prospective epidemiological study PARSIFAL (62). This study compared farmers to non-farmers’ essential markers expressions and the prevalence of asthma; the authors determinate an enhanced expression of genes of the innate immunity such as IRAK-4 and RIPK1 as well as regulatory molecules such as IL-10, TGF-beta, SOCS4, and IRAK-2. (62). Although the correlation of Treg and host defense molecules described is similar to our results, our finding pointed to the epithelium involvement in this persistence immune response.

As was descripted before, several experimental and clinical studies established the correlation between LPS pre-exposure and asthma phenotype abrogation; our study attempted to dissect the changes of a pro-allergic cytokine secreted by the epithelium such as TSLP in this context. In this sense, our results demonstrated that LPS neonatal exposition correlated with the abrogation of TSLP expression in epithelium and in BAL. Therefore, meanwhile allergen-induced TLSP recruits dendritic cells that amplify the Th2 response and reduces Treg cells expansion (27). In the last decades, several neonatal and pregnancy animal models suggested that the transition from the quiescent Th2-polarized fetal immune phenotype towards the more active Th1-pattern of mature adaptive immunity was intrinsically slower in the atopic population, thus increasing the risk of an allergen priming response against environmental antigens. (21, 22, 63–68). Thus, it would be important to future evaluate whether this early proinflammatory stimulus by LPS could cooperate with the progression of this transition.

In our study, LPS diminished the TSLP mRNA basal expression consistent with its intrinsic capacity to counterbalance different pro- allergic action; thus blunted the subsequent overproduction of TSLP in front of OVA exposition. In a model of human bronchial epithelial cell line, Lin et al also demonstrated that LPS pre-treatment could reduce the induction of TSLP mRNA levels by a virus that causes neonatal respiratory disease (69). Interestingly, in this study the basal mRNA levels of different signaling proteins involved in the TSLP overproduction were downregulated only when a repeated LPS- preventive treatment were applied. In a process that, similar to ours results, the authors attributed to the modulation in the expression of innate immunity signaling molecules of the airway epithelial cells to mitigate the allergic response.

The main contribution of our study is highlighting the involvement of bronchioalveolar epithelium in this early microbial protection from allergic disorder, a topic that remains unaddressed. This was demonstrated by the stable changes in the expression of antiallergic proteins as well as host defense factors by CC and the reduction to basal levels of potent epithelial-Th2 mediators, all of which was promoted by neonatal LPS-stimulation that further polarized Treg response in front of allergen exposition. Altogether, our results point to several anti-allergic dynamic mechanisms overlying in the epithelium that could be favored in an adequate epidemiological environment.

